# Metagenomics reveals that Dung Beetles (Coleoptera: Scarabaeinae) broadly feed on Reptile dung and could feed on that of Dinosaurs

**DOI:** 10.1101/2022.12.27.521994

**Authors:** Fernando Lopes, Michele Rossini, Federica Losacco, Giulio Montanaro, Nicole Gunter, Sergei Tarasov

## Abstract

According to traditional views, the evolution of dung beetles (Coleoptera: Scarabaeinae) and their feeding habits are largely attributed to mammal dung. In this paper, we challenge this view and provide evidence that many dung beetle communities are actually associated with the dung of reptiles and birds (= Sauropsida). In turn, this indicates that sauropsid dung may have played a crucial evolutionary role that was previously underestimated. We argue that it is physiologically realistic to consider that coprophagy in dung beetles could have evolved during the Cretaceous in response to the massive amount of dung produced by dinosaurs. Furthermore, we demonstrate that sauropsid dung may be one of the major factors driving the emergence of insular dung beetle communities across the globe. We support our findings with amplicon-metagenomic analyses, trapping experiments, and meta-analysis of the published literature.

## 1 INTRODUCTION

With over 6,000 species, dung beetles or scarabaeines (Coleoptera: Scarabaeinae) mostly feed on mammalian dung, particularly that of herbivores, and serve as primary decomposers of feces on Earth (Nichols et al., 2008; Hanski and Cambefort, 2014) directly affecting ecosystem services and processes like nutrient cycling, soil aeration, and chemical composition (deCastro Arrazola et al., 2022). The dung beetle diversification is suggested to be linked to the radiation of mammals in the Cenozoic (Scholtz et al., 2009; Gunter et al., 2016; O’Leary et al., 2013), while the origin of dung-feeding behavior is proposed to have happened early in dung beetle evolution by shifting from saprophagy to coprophagy (Hanski and Cambefort, 2014). In most cases, dung beetles are found to be associated with the feces of mammals (Stavert et al., 2014). A few other vertebrates whose dung is known to be consumed by dung beetles are birds, reptiles, and amphibians but such records are rare (Fincher et al., 1970; Kabir et al., 1990; Gill, 1991; Stavert et al., 2014; Halffter and Matthews, 1966; Young, 1981).

In the present study, we demonstrate that even though mammalian feces are undeniably important, the role of bird and reptile (= Sauropsida) dung may have been greatly undervalued in the evolution of dung beetles. In Madagascar and Mauritius, we conducted trapping experiments and sequenced the DNA from the dung visited *in natura* by dung beetles using amplicon-metagenomics to identify the dung’s host species. Additionally, we performed a meta-analysis of published literature to retrieve all available records associated with sauropsid dung and to assess their overall proportion. According to our empirical findings and meta-analysis, insular dung beetles often rely on reptile and bird dung as a feeding resource, which was underappreciated previously. Whether feeding on bird and reptile dung was an ancient or a recent trait acquired by the insular communities is not clear and we discuss it below. The most likely explanation for this behavior is that insular dung beetles are generalists (Jones et al., 2012), and that they evolved this trait due to the food deficiency on the islands caused by the absence of native large mammals (Stavert et al., 2014; Ebert et al., 2019; Langton-Myers, 2022).

Our results provide further evidence that physiologically speaking, there is no restriction that the evolution of coprophagy in dung beetles could have been triggered by dinosaur dung. This is in contrast to “the mammalian hypothesis”, broadly accepted by the research community, which suggests the origin of coprophagy along with mammals (Hanski and Cambefort, 2014). Based on available studies, dung beetles are estimated to have originated between 132-40 Mya (Ahrens et al., 2014; Gunter et al., 2016; McKenna et al., 2019; Davis et al., 2017). This period equivocally fits both scenarios regarding the origin of dung-feeding behavior, i.e., dinosaurs vs. mammals. Below, we reevaluate the scenarios in light of our findings. Overall, we provide evidence that sauropsid dung could play a crucial role in the emergence of coprophagy, and that it may be one of the major factors driving the establishment of insular dung beetle communities.

## 2 METHODS

### 2.1 Trapping experiments

In February 2021, we carried out trapping experiments in two localities on Mauritius island: Mount Le Pouce Nature Reserve (−20.1986, 57.5294) and Black River Gorges National Park (−20.3755, 57.4442; −20.4415, 57.47229), where we set up 13 traps baited with chicken dung. In March 2022, we also carried out a trapping experiment in the Kirindy village area, Madagascar (−20.066805, 44.657255) by setting two traps baited with chicken and reptile dung, respectively.

### 2.2 Sampling dung for metagenomics

In March 2022 at Kirindy village area, Madagascar (−20.074773, 44.590188; −20.066805, 44.657255), we collected ~30 fecal samples, of which ~70% contained 136 specimens of dung beetles (Figure 1B). We primarily identified them as belonging to sauropsids. For the identification of the host species, we selected six pellets for amplicon-metagenomic analysis.

**Figure 1.**
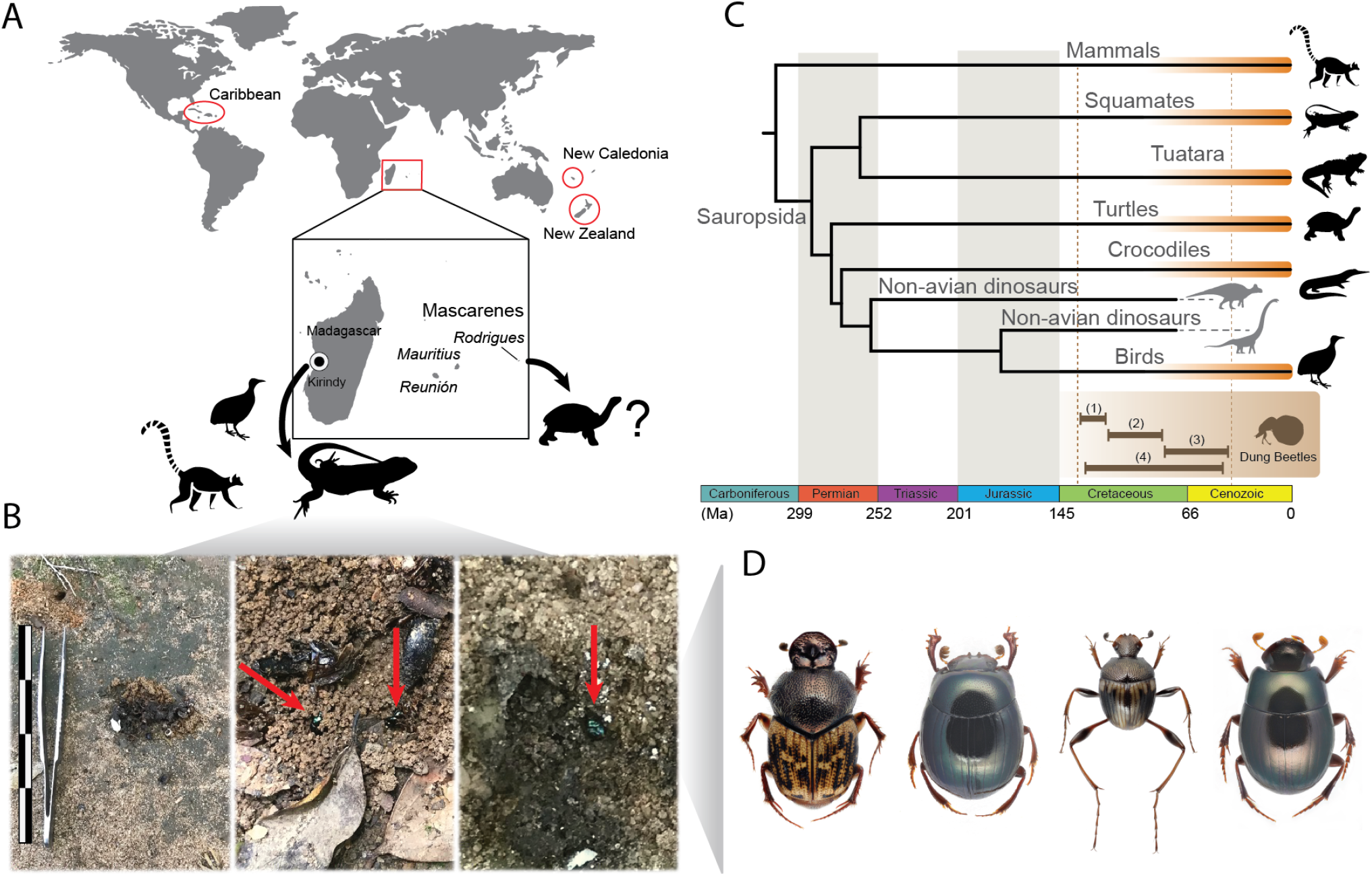
(A) Five island groups with authentic dung beetle communities: Madagascar, Mascarenes, New Caledonia, New Zealand and Caribbean; in the center are the vertebrates whose dung supports dung beetle communities in Madagascar and Mascarenes; the point on Madagascar indicates Kirindy village area. (B) Photos of squamate dung visited by *Helictopleurus* species in Kirindy; the scale bar is 14 cm in length. (C) Phylogenetic tree illustrating which clades of sauropsids and mammals produce suitable dung for scarabaeines (the clades are highlighted in orange); the brown lines on the bottom indicate the ages for the origin of dung beetles according to: (1) Gunter et al. (2016), (2) Ahrens et al. (2014) (3) McKenna et al. (2019), (4) Davis et al. (2017). (D) Dung beetles recorded on sauropsid dung, from left to right: *Helictopleurus infimus* (Madagascar), *Apotolamprus metallicus* (Madagascar), *Nesosisyphus vicinus* (Mauritius), and *Nesovinsonia vinsoni* (Mauritius).

### 2.3 DNA extraction, library preparation, and sequencing

For each sample, we performed DNA extractions using QIAGEN DNeasy PowerSoil Kit and fecal samples (up to 0.25 g) following the manufacturer’s protocol. The purified DNA was quantified using Qubit fluorometer 4.0, high-sensitivity reagents, and 5 μl of DNA extract. The DNA extracts were used for amplicon-metagenomics sequencing using COI universal primers Sauron-878 (5-’GGDRCWGGWTGAACWGTWTAYCCNCC-3’) (Rubbmark et al., 2018) and jgHCO2198 (5’-TAIACYTCIGGRTGICCRAARAAYCA-3’) (Geller et al., 2013) and PCR profile following the protocol of Rubbmark et al. (2018). The PCR products were selected from 2% agarose gel electrophoresis and the same amount from each sample was pooled and tailed with Illumina adapters (Table S1). Libraries were prepared using the NEBNext® Ultra^™^ II DNA Library Prep Kit, checked with Qubit (high-sensitive reagents) and real-time PCR for quantification, and Bioanalyzer for size distribution detection. The amplicon paired-end libraries (PE250) targeting an insert size of 320 bp were sequenced on an Illumina NovaSeq 6000 platform aiming for 30K raw tags (raw reads generated from the opposite end of the same DNA fragment) per sample. Library preparation, quality control, and sequencing were performed at Novogene (Cambridge, UK).

### 2.4 Bioinformatics

Sequenced reads were demultiplexed based on barcode sequences for each sample using Qiime 1.7.0 (Bokulich et al., 2013). Reads with sequencing Phred-score <30 and chimeric reads were filtered out from the downstream pipeline. Cleaned paired-end reads (barcode and primer-free sequences) were merged with FLASH 1.2.7 (Chaisson et al., 2004). The Operational Taxonomic Units (OTUs) clustering was carried out on cleaned and merged paired-end sequences (Effective Tags). To obtain community composition in each fecal sample, OTUs were assembled from sequences with ≥ 97% of similarity using Uparse 10 (Edgar, 2013). The taxonomic annotation was automatically constructed using NCBI BLAST+ 2.13 and the COI database.

From the full annotation table, we extracted and quantified reads assigned to vertebrates (aiming at host identification). We double-checked the OTUs automatically assigned by blasting the sequences manually against the NCBI database with the BLASTn algorithm.

To validate the identification of the hosts, we recovered a Maximum-Likelihood (ML) phylogeny for the COI gene with the most representative OTUs (≥ 94% of the Effective Tags assigned to sauropsids, see Results) from each sample and sequences obtained from GenBank for closely related Madagascan species. Sequences were aligned with MAFFT 7.490 (Katoh and Standley, 2013) and manually checked and trimmed with MEGA X (Kumar et al., 2018). The ML phylogeny of the host species was recovered with IQ-Tree 2.0.3 (Minh et al., 2020), 1000 replicates of ultrafast bootstrap, and SH approximate likelihood ratio test (aLRT).

### 2.5 Meta-analysis of literature

The feeding records data set was assembled using our data and an extensive literature survey on dung beetles (Table S6). In order to test the hypothesis that feeding on sauropsid dung is more likely on islands than on continents, we use a binomial test. In our null hypothesis, finding a sauropsid dung feeder is equally likely on islands and continents. Therefore, the total number of insular records should be proportional to the relative area of islands. This area was calculated by using the data from Sayre et al. (2019) for the islands with an area of over one kilometer squared and by subtracting the area of Greenland (since no dung beetles occur there). The total area of the continental landmasses was calculated as a sum of the areas of Afro-Eurasia, the Americas, and Australia (dung beetles do not occur in Antarctica). The relative area of islands was thus 0.059. According to the null hypothesis, insular dung beetle records should follow the binomial distribution with a probability of 0.059. Our alternative hypothesis proposes that sauropsid dung feeders are more common on islands than on continents. The R script to perform the test is available upon request.

## 3 RESULTS

### 3.1 Dung beetles sampled in sauropsid dung

The trapping experiments carried out in Mauritius clearly showed a high attractiveness of native dung beetles to chicken excrement. The samplings allowed us to recollect four of the five known Scarabaeinae species endemic to the island: *Nesosisyphus regnardi* (Alluaud, 1898) (Mount Le Pouce), *N. pygmaeus* Vinson, 1946 (Mount Ory), *N. vicinus* Vinson, 1939 (Mount Cocotte), and *Nesovinsonia vinsoni* (Paulian, 1939) (Mount Le Pouce). Although the majority of dung beetles were captured using chicken dung, including the only four specimens of the monotypic genus *Nesovinsonia*, some specimens were also attracted by human excrement, and among them the only specimen of *N. pygmaeus* collected during the field trip. It is noteworthy that about 10 specimens of *N. regnardi* were collected using pitfall traps baited with the carrion of the invasive giant snail *Lissachatina fulica* (Bowdich, 1822). The same behavior was also reported for *N. pygmaeus*, which was observed feeding on the excrement of the endemic Mauritian snail *Pachystyla bicolor* (Vinson, 1951).

In Kirindy village, north of the Kirindy National Park, Madagascar, a total of 140 specimens were collected in sauropsid dung. The pitfall trap baited with chicken dung yielded four specimens and four species: *Apotolamprus metallicus* Montreuil, 2008; *Epilissus cuprarius* Fairmaire, 1899; and two unidentified *Arachnodes* Westwood, 1842 species, tentatively related to *A. kelifelyi* Paulian, 1976 and *A. pillula* Paulian, 1976. Instead, reptile dung yielded 136 dung beetle specimens belonging to six species as follows: *Onthophagus pipitzi* Ancey, 1883, *Helictopleurus infimus* (Fairmaire, 1901) and *H. peyrierasi* Paulian & Cambefort, 1991; *A. metallicus; Nanos humeralis* Paulian, 1975; and an unidentified *Arachnodes* species possibly related to *A. kelifelyi* (Figure 1D). During the trapping experiments, *H. infimus* was the most abundant species, followed by *A. metallicus* and *O. pipitzi* (Figure 1D).

### 3.2 Metagenomics: identifying dung hosts

We recovered 349,012 Effective Tags from 399,605 Raw Tags (reads), with an average of 58,170 (SD ± 20.9) Effective Tags per sample. This amount represents 86% (SD ± 8.9) of the raw reads with Phred-score >30 (Table S2). In the process of constructing OTUs, summary information from different samples was collected, such as Effective Tags, low-frequency Tags, and Tags’ annotation. The summary is shown in Figure S1.

We found a relatively high alpha (Figure S2) and beta diversity (Figures S3, S4), with the majority of the reads belonging to microorganisms like bacteria and fungi (Figure S2). This high diversity of microorganisms was expected since the source of the DNA was organic material (excrement) in different stages of decomposition. However, from the total number of Effective Tags generated, 40,743 were assigned to vertebrates. Of this number, 99.95% were assigned to species of reptiles (Figure 2A), while the remaining 0.05% were assigned to mammals and therefore considered to be contamination (Tables S3, S4, and Figure S5). Under this number and among the six fecal samples, the proportion of reads assigned to the same species of reptile ranged from 94.2 to 100%, indicating that the most representative OTUs probably represent the host species (Figure 2A). We found no traces of dung beetles (Scarabaeinae) among the sequences identified as Coleoptera (Table S5).

**Figure 2.**
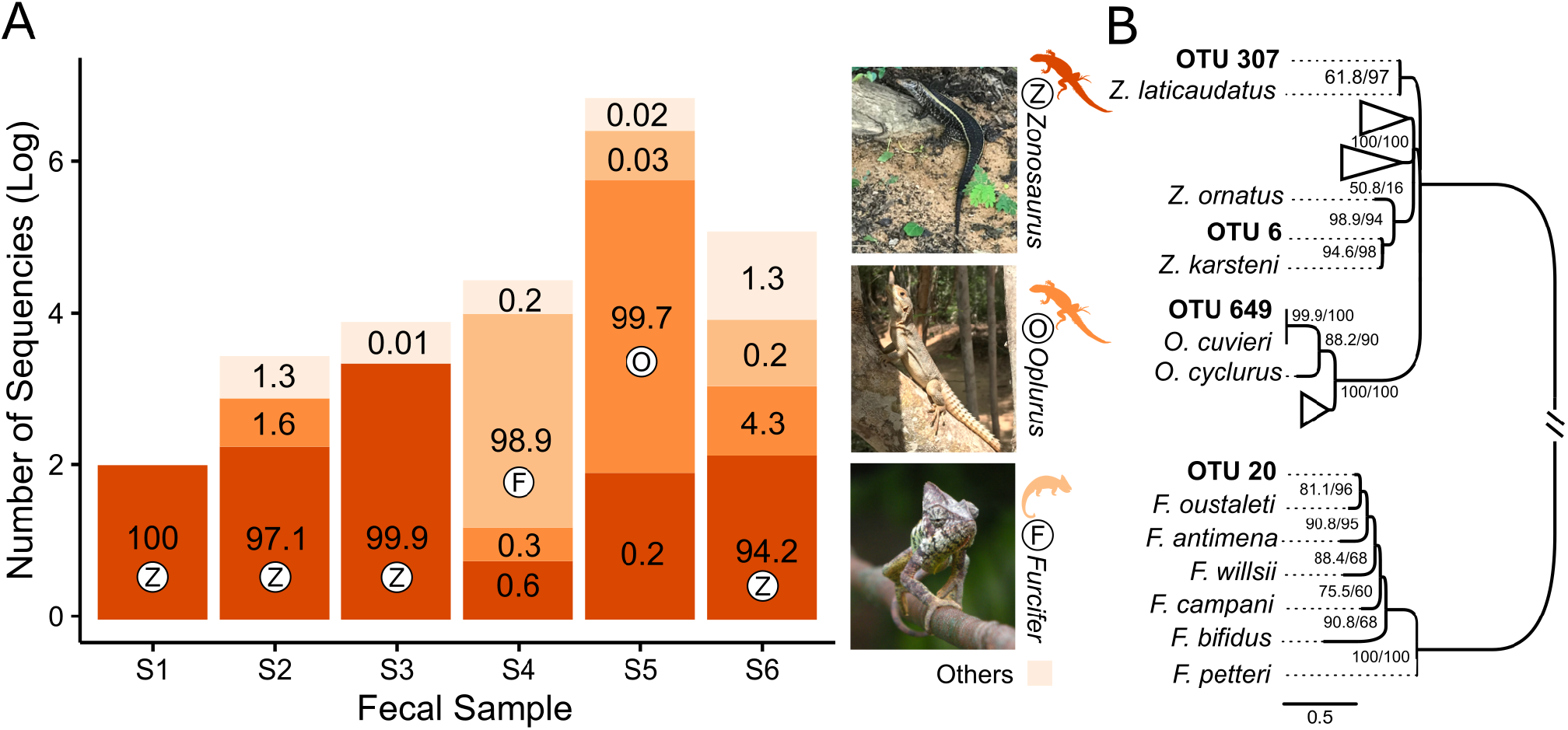
Results of the amplicon-metagenomic analysis. (A) The proportion of Effective Tags (log-scale) within each fecal sample (S1-S6); numbers in the bars show relative abundance (%) of reads; colors and letters (Z, F, O) represent squamate genera shown on the right; in each fecal sample, the genus with the highest proportion of Effective Tags is marked with the respective letter. (B) Maximum-Likelihood tree, based on COI region, that shows the phylogenetic position of squamate OTUs from the fecal samples; the node numbers are the ultra-fast bootstrap support and SH-aLRT values, respectively; full phylogeny is available in Figure S6. The picture of *Furcifer oustaleti* (male), Lake Ravelobe, by Frank Vassen is licensed under CC BY 2.0.

The phylogeny inferred using the COI gene, OTUs, and sequences obtained from GenBank (Figures 2B and S6) validated the OTU assignment and allowed the identification of the hosts at the species level. The phylogeny indicated four host species belonging to three families, respectively: fecal samples S1, S2, and S6 clustered with *Zonosaurus laticaudatus* (Grandidier, 1869) and sample S3 with *Z. karsteni* (Grandidier, 1869) (Iguanidae); sample S4 with *Furcifer oustaleti* (Mocquard, 1894) (Chamaleonidae); and sample S5 with *Oplurus cuvieri* Gray, 1831 (Opluridae) (Figures 2B and S6). The distribution maps available on the IUCN Red List of Threatened Species (iucnredlist.org) and iNaturalist (inaturalist.org) databases confirm that all species potentially occur at Kirindy National Park and surrounding areas.

### 3.3 Meta-analysis of literature

Our survey found 79 records reporting the association between scarabaeines and sauropsid dung (Table S6). Sixteen records were insular (Madagascar, Mauritius, New Zealand, Puerto Rico), while the other 63 were continental. The continental records cover the Americas, Australia, Asia, and Africa. Dung from all major Sauropsida clades (Squamates, Tuataras, Turtles, Crocodiles, and Birds) was found to attract dung beetles (Figure 1C).

The one-sided binomial test rejects the null hypothesis that finding a sauropsid dung feeder is equally likely on islands and continents (p-value = 0.00001). Contrary to that, our findings support the alternative hypothesis that sauropsid dung feeders are 3.4 times more likely to be found on islands.

## 4 DISCUSSION

### 4.1 Insular dung beetles communities

According to our binomial test of the feeding records published from 1894 to the present, it is 3.4 times more likely to find dung beetles feeding on sauropsid dung on the island than on the continent. There are five island groups (Figure 1A) that harbor authentic scarabaeine communities with endemic genera: Madagascar, Mascarenes (Mauritius, Réunion, Rodrigues), New Caledonia, New Zealand and Caribbean (Davis et al., 2002). Our findings suggest that almost all of them possess a significant proportion of scarabaeines associated with sauropsid dung. With the use of COI amplicon-metagenomics, we were able to identify three lizard and one chameleon species whose dung was consumed by at least five different species of dung beetles in Madagascar (Figure 1 and Table S6). In addition, five other scarabaeine species were recorded by us in earlier studies as visitors and potential feeders of sauropsid dung in Madagascar and Mauritius (Table S6). The widespread consumption of sauropsid dung has also been documented in New Zealand (Stavert et al., 2014), and hypothesized to have existed in the Caribbean, where only a single record has been found so far (Halffter and Matthews, 1966; Matthews, 1965). A lack of research into the feeding behavior of scarabaeines from New Caledonia precludes any conclusions regarding their diet.

Recent discoveries have indicated that the Mascarene islands used to harbor a significant diversity of dung beetles, which deserves special attention. The Mascarene archipelago consists of three main islands: Mauritius, Réunion, and Rodrigues. The latter is a small volcanic island located ~ 1,500 km east of Madagascar. Extant endemic dung beetles occur only on Mauritius: one species of *Nesovinsonia* Martínez & Pereira, 1958, and four species of *Nesosisyphus* Vinson, 1946. One extinct endemic species from the genus *Epactoides* Olsouffief, 1947 was recently found on Réunion (Rossini et al., 2021). It is noteworthy that large deposits of extinct dung beetles were recently discovered on Mauritius and Rodrigues [Nick Porch personal communication, see also Rijsdijk et al. (2009)]. This indicates that the scarabaeine diversity on Mascarenes was greater in the past than it is today. Due to the lack of native mammals on these islands, other than bats, it is plausible that such a diversity of dung beetles was established and maintained by the dung of giant tortoises and/or dodos, which were prevalent on the islands prior to human arrival (Rijsdijk et al., 2009; Galetti et al., 2018). We discuss below why we rule out alternate feeding options in this paper.

Most species of dung beetles that have been recorded on islands are generalists since they have been observed to consume other food sources as well (Jones et al., 2012; Ebert et al., 2019; Langton-Myers, 2022). Only two species, *Copris gopheri* Hubbard, 1894 and *Onthophagus polyphemi* Hubbard, 1894, are known to feed exclusively on the dung of the Gopher tortoise *(Gopherus polyphemus)* from the USA (Hubbard, 1894; Young and Goff, 1939; Howden and Cartwright, 1963). These facts indicate that dung beetle communities on islands are strongly associated with sauropsid dung, but that they are not solely dependent on it. The authentic insular communities emerged as a result of colonization events from neighboring landmasses. For example, the Madagascan lineages originated from Africa approximately 38-8 Mya (Miraldo et al., 2011; Rossini et al., 2022). The endemic clades found in New Caledonia and New Zealand dispersed from Australia at 50-55 Mya (Gunter et al., 2016). According to this evidence, it is reasonable to assume that the first colonizers evolved the generalist diet, which included feeding on sauropsid dung, as a way to circumvent food shortages caused by the lack of large mammals on the islands (Cupello et al., 2020). Nevertheless, a conclusion of this nature requires further investigation in the phylogenetic framework in order to reconstruct ancestral diets and refine colonization times.

Dung beetles evolved adaptations to many diets other than dung, including carrion, fungi, and detritus [reviewed in Scholtz et al. (2009)]. Hence, it is important to discuss why we rule out a pivotal role for at least some of those adaptations in maintaining insular communities. For example, islands have large populations of bats that produce large amounts of guano. Do insular scarabaeines feed on guano? In total, we found only nine records (Table S6) reporting an association between bat guano and dung beetles, and they all belong to continental environments. Thus, despite the abundance of bats on islands, there is no evidence that local dung beetles have ever fed on bat dung.

As regards other sources such as plant debris, carrion, and fungi, it is difficult to quantify their role on islands compared to continents based on available data. For example, many investigations report that dung beetles in New Zealand are saprophagous (Ecroyd, 1996; Hodge et al., 2010; Jones et al., 2012). Because of the lack of rigorous data, it is challenging to estimate the proportion and distribution of saprophagy in continental scarabaeines, which precludes any comparative analysis. Therefore, in assessing the different diets, we rely primarily on the present results due to their statistical and empirical support, and these results indicate an increased proportion of sauropsid dung feeders on islands. Yet, we must acknowledge that further research on diet distribution is needed, which may well reveal that sauropsid dung, saprophagy, and carrion-feeding, are all fundamental to the existence of the insular dung beetles.

### 4.2 Dung beetles and dinosaurs

Currently, the only direct evidence of an association between dung beetles and dinosaurs comes from the coprolite of a Cretaceous dinosaur. The burrows in this coprolite have been attributed to dung beetle activity (Chin and Gill, 1996). However, there is no evidence to support a reliable connection between those burrows and a dung beetle (Tarasov et al., 2016). Furthermore, additional evidence has questioned the association of scarabaeines with dinosaurs. It has been suggested that dinosaurs, like birds, produced feces containing ammonia, uric acid, and other chemical components that were not attractive to dung beetles (Arillo and Ortuño, 2008). Additionally, dinosaurs’ metabolism may prevent them from producing sufficient amounts of dung to support dung beetle communities (Scholtz et al., 2009). It is important to stress that these arguments were primarily based on the fact that the majority of extant dung beetles do not feed on the dung of reptiles. The purpose of this paper was to demonstrate the opposite: at least some dung beetle communities do broadly feed on the feces of sauropsids. In fact, we found that all major clades of extant sauropsids, including squamates, tuataras, turtles, crocodiles, and birds, produce dung suitable for dung beetles (Figure 1C). Therefore, it is logical to assume that past scarabaeines were physiologically capable of consuming dinosaur dung, just as extant scarabaeines consume that of sauropsids.

According to available studies, the timing of the origin of the dung beetles is unclear due to uninformative fossil records (Tarasov et al., 2016) and the lack of a credible time tree. Published estimates place the emergence of scarabaeines between 132 and 40 Mya (Ahrens et al., 2014; Gunter et al., 2016; McKenna et al., 2019; Davis et al., 2017). Considering that these estimates overlap the K-Pg boundary, it is also possible that dinosaur dung could have triggered the evolution of coprophagy in dung beetles if the latter coexisted with the former, a hypothesis explored in depth by Gunter et al. (2016). This study estimated the origin of dung feeding in dung beetles in the Upper Cretaceous, a timeline that corresponded with diverse and large sauropod fauna and the rise of the angiosperms. Gunter et al. (2016) hypothesized that a shift in dinosaur diet that included incidental ingestion of the more nutritious and less fibrous angiosperm foliage, provided a palatable dung source with appropriate particle size and moisture content that allowed for the key innovation of dung feeding. Further justification for the importance of sauropsids to the evolution of dung beetles was based on generalized traits of extant dung beetles, that is, very few extant scarabaeines feed on small, dry dung pellets of insectivorous mammals that would have been ecologically similar to the Upper Cretaceous mammalian fauna (Gunter et al., 2016). In the absence of fossils from this time period and therefore reliable ages estimates, such hypotheses remain speculative, although we anticipate that future phylogenetic studies with refined ages will be necessary to answer the question regarding the relationship between dinosaurs and dung beetles.

## 5 CONCLUSION

This study demonstrates the importance of considering atypical dung beetle communities, such as those from islands, to gain a deeper understanding of the evolution of dung beetles. While mammals have undoubtedly played a significant role in the diversification of dung beetles, the contribution of sauropsids should not be overlooked. The extent of how widespread feeding on reptile dung or bird droppings is in extant dung beetles remains unknown, and it is likely there is a bias in studies conducted with alternative resources of non-mammalian dung feeding from areas that lack a diverse fauna of these vertebrates. For example, Australian native dung beetles are widely reported to be generalists (Matthews, 1974; Gunter et al., 2019), and as such multiple bait types are commonly used for specimen collection. Regardless, this literature review identified reports of sauropsid dung feeding in almost all biogeographic realms where dung beetles occur, with the exception of Indomalayan, including countries with diverse mammal faunas. On the basis of this manuscript, we recommend that future studies on dung feeding also consider sauropsid dung as a resource. For example, the use of non-mammalian dung options in feeding ecology studies, the use of broad vertebrate primers, or database screening beyond Mammalia in molecular gut content analyses will provide important positive and negative association data that will help disentangle the extent of generalist feeding in dung beetles. We expect that generalist feeding ecology is more widespread than currently reported, and this knowledge is essential to our understanding of the evolutionary history of dung beetles, as well as predicting the impact of mammal decline on them.

## AUTHOR CONTRIBUTIONS

ST and MR collected samples. ST and FLOP designed the study. Data analyses were performed by FLOP with the contribution of ST, MR, and NG. Morphological identification of the dung beetles was performed by MR with the support of GM and FLOS. The initial manuscript draft was written by FLOP, MR, NG, and ST. All authors edited, contributed to the interpretation of results, and approved the final version of the manuscript.

## ACKNOWLEDGMENTS

We are very thankful to the Malagasy Institut pour la Conservation des Ecosystèmes Tropicaux (MICET) for the support in the acquisition of permits and logistic support during the fieldwork; The Mauritian National Parks and Conservation Service (NPCS) for providing permits to collect in Mauritius; Vincent Florens (University of Mauritius) and Owen Griffiths (La Vanille Nature Park, Mauritius) for the logistic support and continuous help during our 2021 expedition to Mauritius; Philippe Moretto for sharing his extensive bibliography on scarab beetles. All people in the Tarasov Lab for their general suggestions and discussions. We are also thankful to the staff of the DNA Lab of the Finnish Museum of Natural History for the wet-lab support.

## DATA AVAILABILITY STATEMENT

Amplicon raw reads for this study can be retrieved from NCBI Sequence Read Archive (SRA) database under the BioProject PRJNA889602. Assembled OTUs, alignments, the complete annotation table, and the Table S6 are accessible at (DOI: 10.17605/OSF.IO/JH3AM).

All biological samples were collected, transported, and exported under the following permits. N°443/21/MEDD/SG/DGGE/DAPRNE/SCBE.Re; N° 002-08/MEEFT/SG/DGEF/DVRNE/SADG; 75-22/MEDD/SG/DRDD.ALM/CIREF MOR; 102-22/MEDD/SG/DRDD.ALM (Madagascar). NP 46/3V5 (Mauritius).

