## Supplementary Tables and Figures for "Metagenomics reveals that Dung Beetles (Coleoptera: Scarabaeinae) broadly feed on Reptile dung and could feed on that of Dinosaurs"

### Supplementary Material

#### 1 SUPPLEMENTARY MATERIAL

##### 1.1 Morphological Identification of Dung Beetles

The morphological examination of the dung beetles collected in the fecal samples was carried out under a Leica S9D stereomicroscope. Photographs were taken with a Canon EOS 5D camera and a Canon MP-E 65 mm, f/2.8, 1–5× macro lens, using the Cognisys Stackshot automated system. Images were subsequently enhanced in Adobe Photoshop, and figures were assembled and edited in Illustrator CC. Specimen identification was carried out by morphological comparison with material deposited in the Coleoptera collection of the Finnish Museum of Natural History (LUOMUS), Helsinki (FMNH), as well as consultation of the main taxonomic literature (Paulian, 1975; Paulian and Cambefort, 1991; Montreuil,2008; Rossini et al., 2021a).

#### 2 SUPPLEMENTARY TABLES

**Table S1.** Barcode and sequences of the primers Sauron-878 and jgHCO2198 used for each sample.

| ID | Barcode Sequence | Primer Sequence |
|---|---|---|
| S1 | CAATGGAA,CATACCAA | GGDRCWGGWTGAACWGTWTAYCCNCC,TAIACYTCIGGRTGICCRAARAAYC |
| S2 | CAATGGAA,CCAGTTCA | GGDRCWGGWTGAACWGTWTAYCCNCC,TAIACYTCIGGRTGICCRAARAAYC |
| S3 | CAGCGTTA,CATACCAA | GGDRCWGGWTGAACWGTWTAYCCNCC,TAIACYTCIGGRTGICCRAARAAYC |
| S4 | CACTTCGA,CCGAAGTA | GGDRCWGGWTGAACWGTWTAYCCNCC,TAIACYTCIGGRTGICCRAARAAYC |
| S5 | CACTTCGA,CCAGTTCA | GGDRCWGGWTGAACWGTWTAYCCNCC,TAIACYTCIGGRTGICCRAARAAYC |
| S6 | CACTTCGA,CATACCAA | GGDRCWGGWTGAACWGTWTAYCCNCC,TAIACYTCIGGRTGICCRAARAAYC |

**Table S2.** Summarization of the data processed for each sample.

| ID | Raw PE | Raw Tags | Clean Tags | Effective Tags | Length (bp) | Q30 | GC (%) | Effective (%) |
|---|---|---|---|---|---|---|---|---|
| S1 | 62,314 | 58,168 | 58,013 | 58,013 | 319 | 97,12 | 30,36 | 93,1 |
| S2 | 76,983 | 72,081 | 71,729 | 71,729 | 314 | 97,38 | 30,5 | 93,18 |
| S3 | 39,174 | 30,757 | 28,286 | 28,285 | 311 | 96,86 | 42,11 | 72,2 |
| S4 | 99,192 | 88,022 | 87,615 | 87,615 | 314 | 95,58 | 33,96 | 88,33 |
| S5 | 67,111 | 61,928 | 61,028 | 61,028 | 314 | 97,22 | 38,49 | 90,94 |
| S6 | 54,831 | 46,368 | 42,342 | 42,342 | 315 | 97,07 | 37,06 | 77,22 |

**Table S3.** Total amount of reads assigned to reptiles or mammals in each sample. The relative abundance of reads for each taxon is also demonstrated in the last row of the table.

| ID | Reptile | Mammal |
|---|---|---|
| S1 | 166 | 0 |
| S2 | 310 | 4 |
| S3 | 17806 | 4 |
| S4 | 1197 | 3 |
| S5 | 15963 | 3 |
| S6 | 5279 | 8 |
| **Total** | **40721/99.95%** | **22/0.05%** |

**Table S4.** Total amount and the relative abundance (%) of reads assigned to each sample and genus.

| ID | Zonosaurus | Oplurus | Furcifer | Homo | Bos | Pteromys | Bad ID |
|---|---|---|---|---|---|---|---|
| **S1** | 166/100 | 0/0 | 0/0 | 0/0 | 0/0 | 0/0 | 0/0 |
| **S2** | 304/97.12 | 5/1.60 | 0/0 | 2/0.64 | 2/0.64 | 0/0 | 0/0 |
| **S3** | 17806/99.98 | 0/0 | 0/0 | 0/0 | 4/0.02 | 0/0 | 0/0 |
| **S4** | 7/0.58 | 3/0.25 | 1186/98.92 | 3/0.25 | 0/0 | 0/0 | 0/0 |
| **S5** | 38/0.24 | 15920/99.71 | 5/0.03 | 3/0.02 | 0/0 | 0/0 | 0/0 |
| **S6** | 5042/94.21 | 228/4.26 | 9/0.17 | 3/0.06 | 2/0.04 | 3/0.06 | 65/1.21 |

**Table S5.** Identification of Coleoptera. The number of Effective Tags is in decreasing order within each sample.

| ID | OTU | Effective Tags | Order | Family | Subfamily | Genus |
|---|---|---|---|---|---|---|
| S1 | OTU 12 | 2090 | Coleoptera | Staphylinidae | Oxytelinae | Carpelimus |
| S1 | OTU 648 | 1415 | Coleoptera | Staphylinidae | Oxytelinae | Carpelimus |
| S1 | OTU 62 | 42 | Coleoptera | Scarabaeidae | Dynastinae | Mimeoma |
| S1 | OTU 16 | 39 | Coleoptera | Cerambycidae | Lamiinae | Eutetrapha |
| S1 | OTU 434 | 5 | Coleoptera | Staphylinidae | Oxytelinae | Carpelimus |
| S1 | OTU 275 | 5 | Coleoptera | Staphylinidae | Oxytelinae | Carpelimus |
| S1 | OTU 966 | 3 | Coleoptera | Staphylinidae | Oxytelinae | Carpelimus |
| S1 | OTU 770 | 2 | Coleoptera | Elateridae | Negastriinae | Zorochros |
| S1 | OTU 163 | 2 | Coleoptera | Scarabaeidae | Dynastinae | Mimeoma |
| S1 | OTU 191 | 1 | Coleoptera | Scarabaeidae | Cetoniinae | Glycyphana |
| S2 | OTU 16 | 2364 | Coleoptera | Cerambycidae | Lamiinae | Eutetrapha |
| S2 | OTU 648 | 22 | Coleoptera | Staphylinidae | Oxytelinae | Carpelimus |
| S2 | OTU 12 | 15 | Coleoptera | Staphylinidae | Oxytelinae | Carpelimus |
| S2 | OTU 62 | 3 | Coleoptera | Scarabaeidae | Dynastinae | Mimeoma |
| S2 | OTU 1439 | 2 | Coleoptera | Curculionidae | Entiminae | Polydrusus |
| S3 | OTU 191 | 22 | Coleoptera | Scarabaeidae | Cetoniinae | Glycyphana |
| S3 | OTU 163 | 18 | Coleoptera | Scarabaeidae | Dynastinae | Mimeoma |
| S3 | OTU 162 | 9 | Coleoptera | Scarabaeidae | Dynastinae | Mimeoma |
| S3 | OTU 648 | 6 | Coleoptera | Staphylinidae | Oxytelinae | Carpelimus |
| S3 | OTU 12 | 4 | Coleoptera | Staphylinidae | Oxytelinae | Carpelimus |
| S3 | OTU 182 | 1 | Coleoptera | Scarabaeidae | Dynastinae | Mimeoma |
| S4 | OTU 182 | 2 | Coleoptera | Scarabaeidae | Dynastinae | Mimeoma |
| S5 | OTU 16 | 4 | Coleoptera | Cerambycidae | Lamiinae | Eutetrapha |
| S5 | OTU 62 | 3 | Coleoptera | Scarabaeidae | Dynastinae | Mimeoma |
| S5 | OTU 450 | 1 | Coleoptera | Scarabaeidae | Dynastinae | Mimeoma |
| S6 | OTU 176 | 27 | Coleoptera | Hydrophilidae | | Cymbiodyta |
| S6 | OTU 182 | 10 | Coleoptera | Scarabaeidae | Dynastinae | Mimeoma |
| S6 | OTU 162 | 8 | Coleoptera | Scarabaeidae | Dynastinae | Mimeoma |
| S6 | OTU 450 | 7 | Coleoptera | Scarabaeidae | Dynastinae | Mimeoma |
| S6 | OTU 648 | 6 | Coleoptera | Staphylinidae | Oxytelinae | Carpelimus |
| S6 | OTU 163 | 5 | Coleoptera | Scarabaeidae | Dynastinae | Mimeoma |
| S6 | OTU 12 | 5 | Coleoptera | Staphylinidae | Oxytelinae | Carpelimus |
| S6 | OTU 62 | 4 | Coleoptera | Scarabaeidae | Dynastinae | Mimeoma |
| S6 | OTU 966 | 2 | Coleoptera | Staphylinidae | Oxytelinae | Carpelimus |
| S6 | OTU 1356 | 2 | Coleoptera | Chrysomelidae | Chrysomelinae | Hydrothassa |

#### 3 SUPPLEMENTARY FIGURES

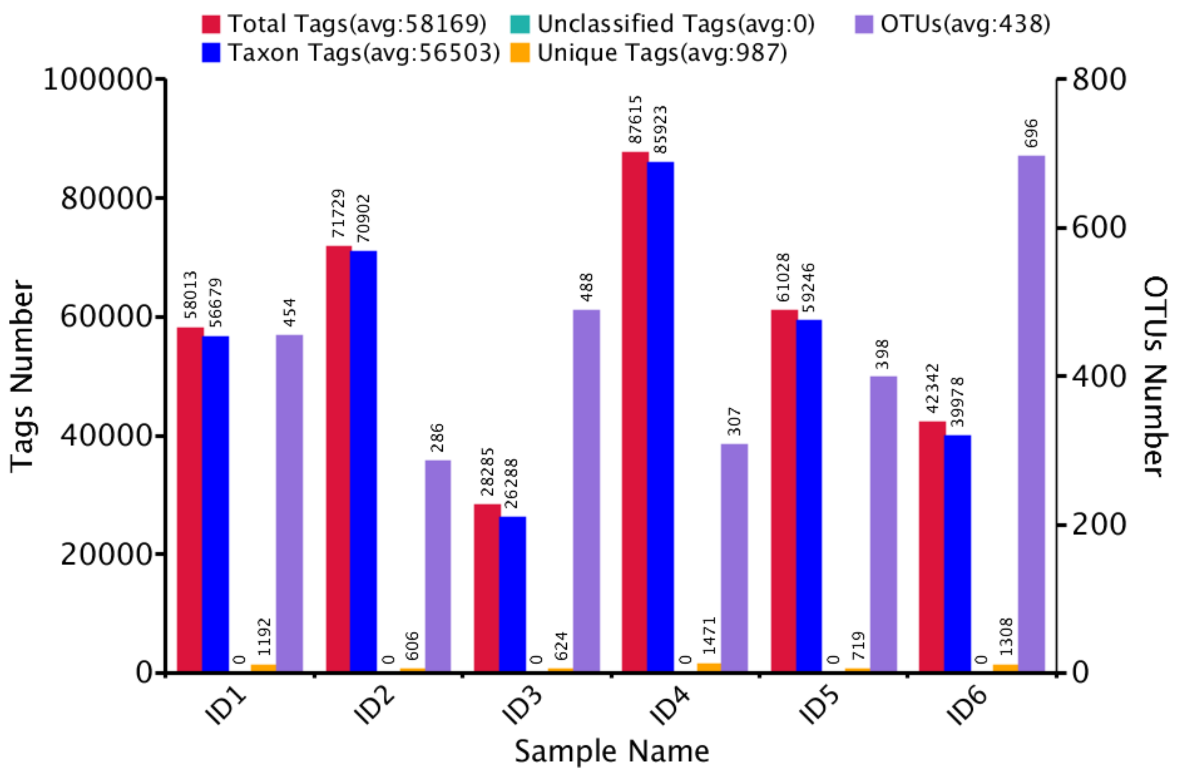


**Figure S1.** Total number of Tags and OTUs per sample

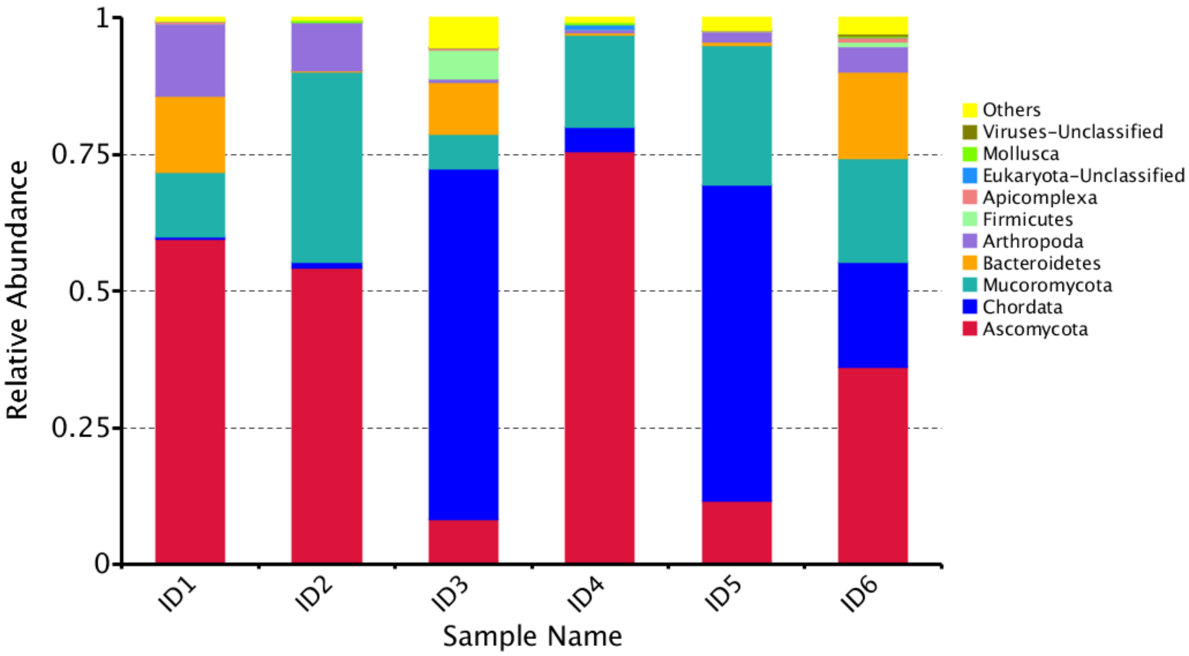


**Figure S2.** Top 10 taxa relative abundance per phylum per sample

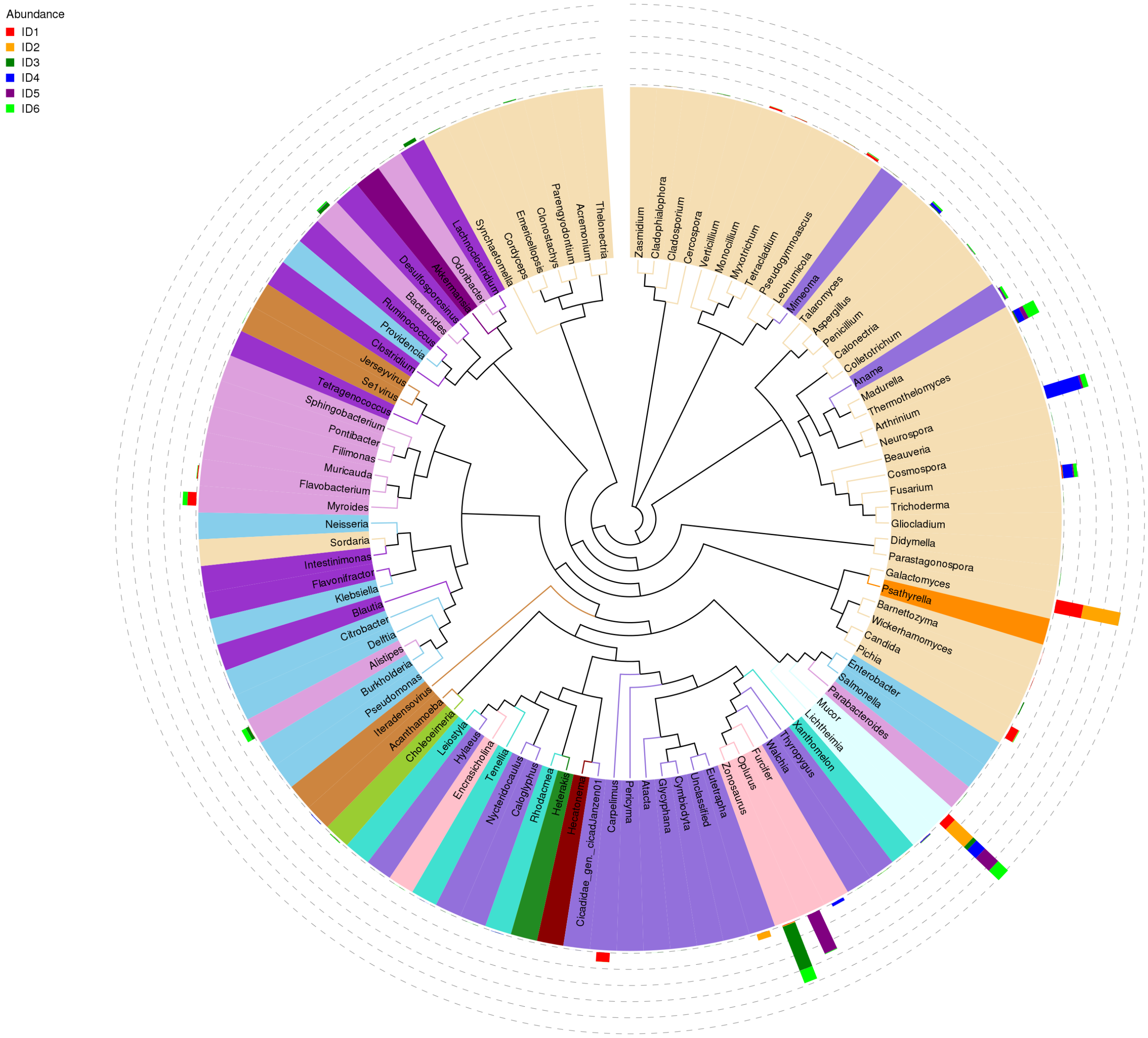


**Figure S3.** Phylogenetic tree (UPGMA) showing clusters at genus level

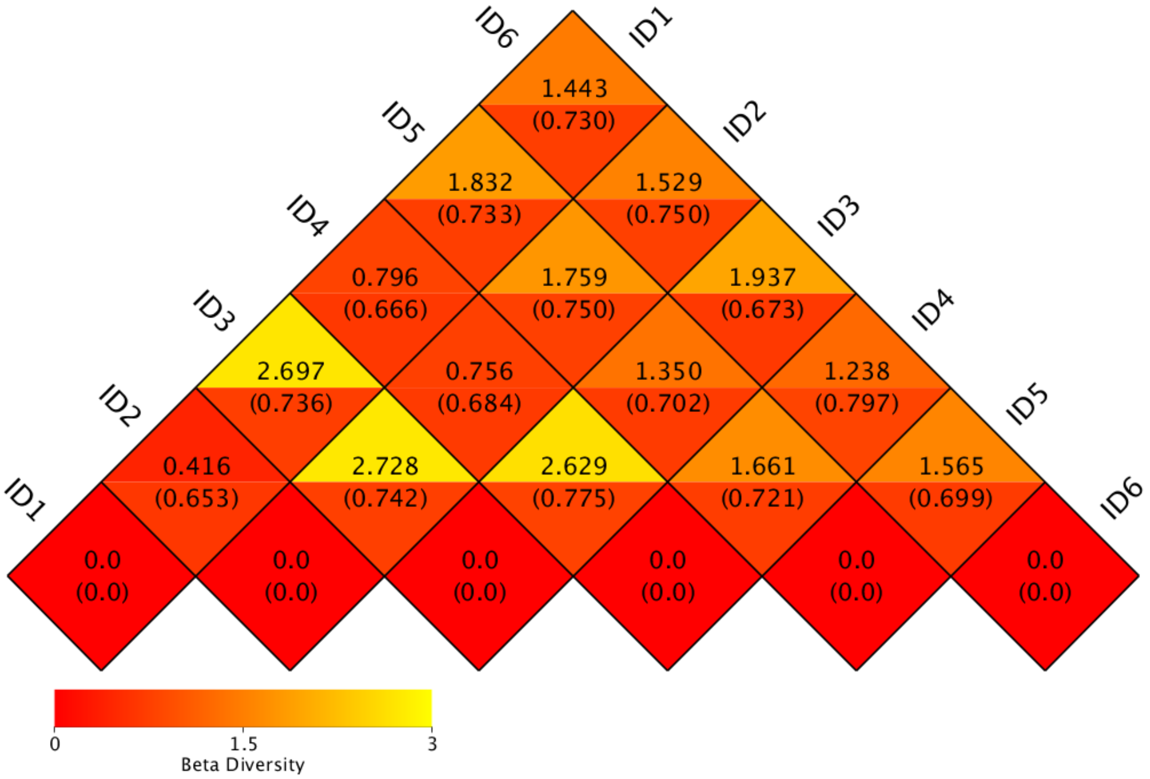


**Figure S4.** Beta diversity. Weighted Unifrac distance and Unweighted Unifrac distance were selected to measure the dissimilarity coefficient between pairwise samples. Each grid represents a pairwise dissimilarity coefficient between pairwise samples, in which the Weighted Unifrac distance is displayed above and the Unweighted Unifrac distance conversely. The smaller the value, the smaller the difference between the two samples in terms of species diversity.

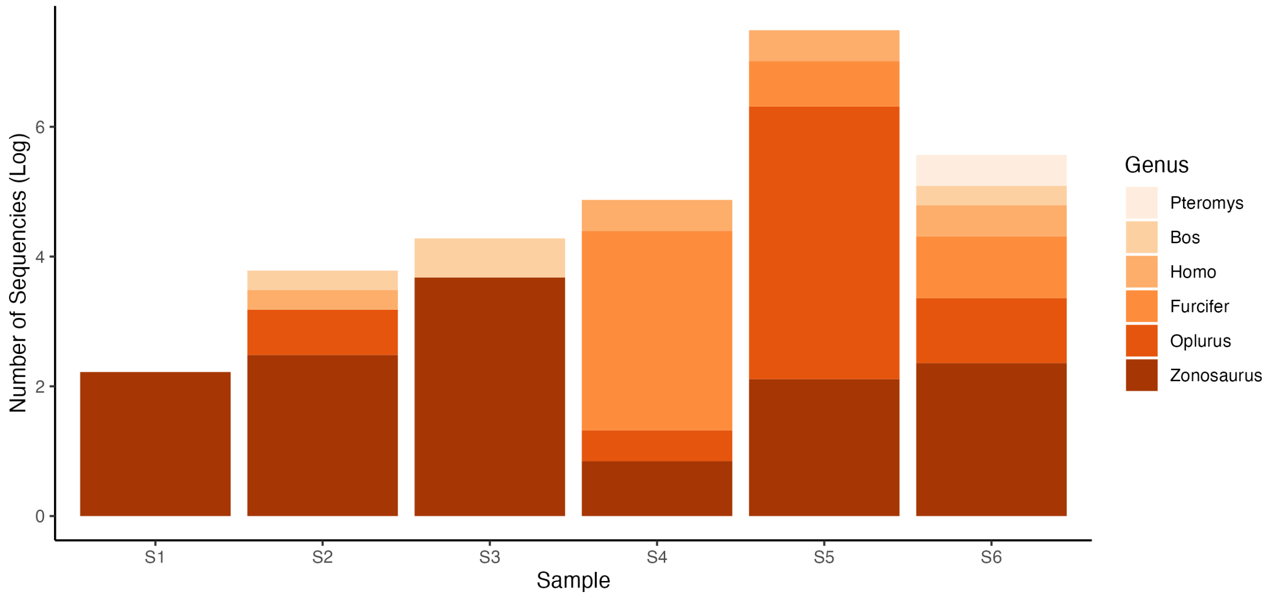


**Figure S5.** Number of sequences per genus in each sample. The number of sequences was log-normalized to highlight contamination.

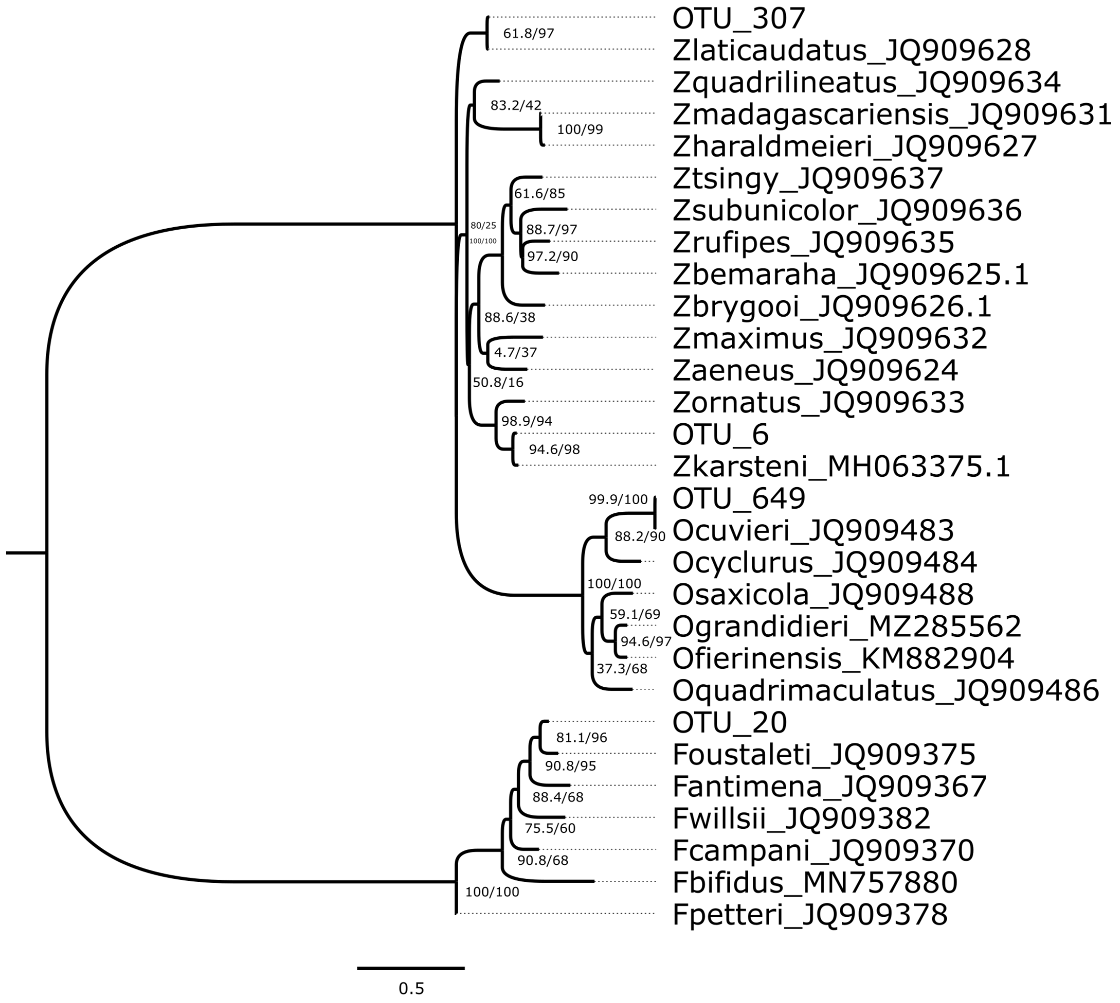


**Figure S6.** Maximum-Likelihood tree of the COI region for the most representative OTUs of each sample and sequences obtained from GenBank. The codes showed with species names are accession numbers from Genbank and the numbers close to the nodes are SH ratio test and bootstrap values, respectively.
